# The Genomic Prehistory of the Indigenous People of Uruguay

**DOI:** 10.1101/2021.11.11.468260

**Authors:** John Lindo, Rosseirys De La Rosa, Andre Luiz Campelo dos Santos, Mónica Sans, Michael DeGiorgio, Gonzalo Figueiro

## Abstract

The prehistory of the people of Uruguay is greatly complicated by the dramatic and severe effects of European contact, as with most of the Americas. After the series of military campaigns that exterminated the last remnants of nomadic peoples, Uruguayan official history masked and diluted the former indigenous ethnic diversity into the narrative of a singular people that all but died out. Here we present the first whole genome sequences of the Indigenous people of the region before the arrival of Europeans, from an archaeological site in eastern Uruguay that dates from 2,000 years before present. We find a surprising connection to ancient individuals from Panama and eastern Brazil, but not to modern Amazonians. This result may be indicative of a distinct migration route into South America that may have occurred along the Atlantic coast. We also find a distinct ancestry previously undetected in South America. Though this work begins to piece together some of the demographic nuance of the region, the sequencing of ancient individuals from across Uruguay is needed to better understand the ancient prehistory and genetic diversity that existed before European contact, thereby helping to rebuild the history of the indigenous population of what is now Uruguay.

## Introduction

Historically, Uruguayan identity has been marked by the extermination of the indigenous populations found in the territory at the time of European contact in the 16th century and up until the 19th century. The extermination was carried out through a series of military campaigns of which the massacre at Salsipuedes creek in 1831 is considered the culminating event.(1) The target of the Salsipuedes campaign was the ethnic group known as the Charrúa, which at the time was the term employed for the remnants of various hunter-gatherer groups in the recently independent territory of Uruguay. Subsequently, it was held that in sharp contrast to all other South American countries, Uruguay lacked indigenous populations, an idea still widely accepted.

The research presented here aims to elucidate the genomic prehistory of the Indigenous people of Uruguay by presenting low-coverage whole genomes from the CH2D01-A archeological site in Rocha, Uruguay, which date from ∼1,450 to ∼668 years before present (BP; Table 1). This represents the first ancient genomic DNA from the region and presents a starting point to examine the evolutionary history of the Indigenous people of Uruguay and their diversity from a genomic perspective.

**Table 1.** Ancient samples sequenced in this study from the archaeological site CH2D01-A inRocha, Uruguay. ^*^Radiocarbon dating was conducted at the University of Arizona AMS Laboratory.^‡^Contamination was estimated from mtDNA alignments using Schmutzi(2).

| Sample | <sup>14</sup>C Age* | Age at death | mtDNA Haplogroup | Contamination<sup>‡</sup> | Endogenous DNA | Average Sequencing Depth | Genome Coverage |
| --- | --- | --- | --- | --- | --- | --- | --- |
| CH-13 | 668 ±22 BP | 45 | C1d3 | 1%,<br>95% CI[0-2%] | 3.4% | 10.7 | 0.137 |
| CH-19B | 1450±70 BP | 18 | C1c | 5%,<br>95% CI[4-6%] | 5.1% | 7.14 | 0.344 |

## Results

To assess the relationship of the ancient Uruguay samples with global and regional populations, both ancient and modern, we merged the dataset with samples from the Simons Genomes Diversity Project (3)and ancient whole genomes from the Americas(4–8). To prevent a batch effect from the method used to call the ancient Uruguay genotypes, the ancient reference BAM files were also called with the ancient DNA caller ARIADNA(9) (*see* Extended Methods). To maximize the number of overlapping sites for the various population genetic analyses, only whole genomes were chosen for comparison (Fig 1A).

**Figure 1.**
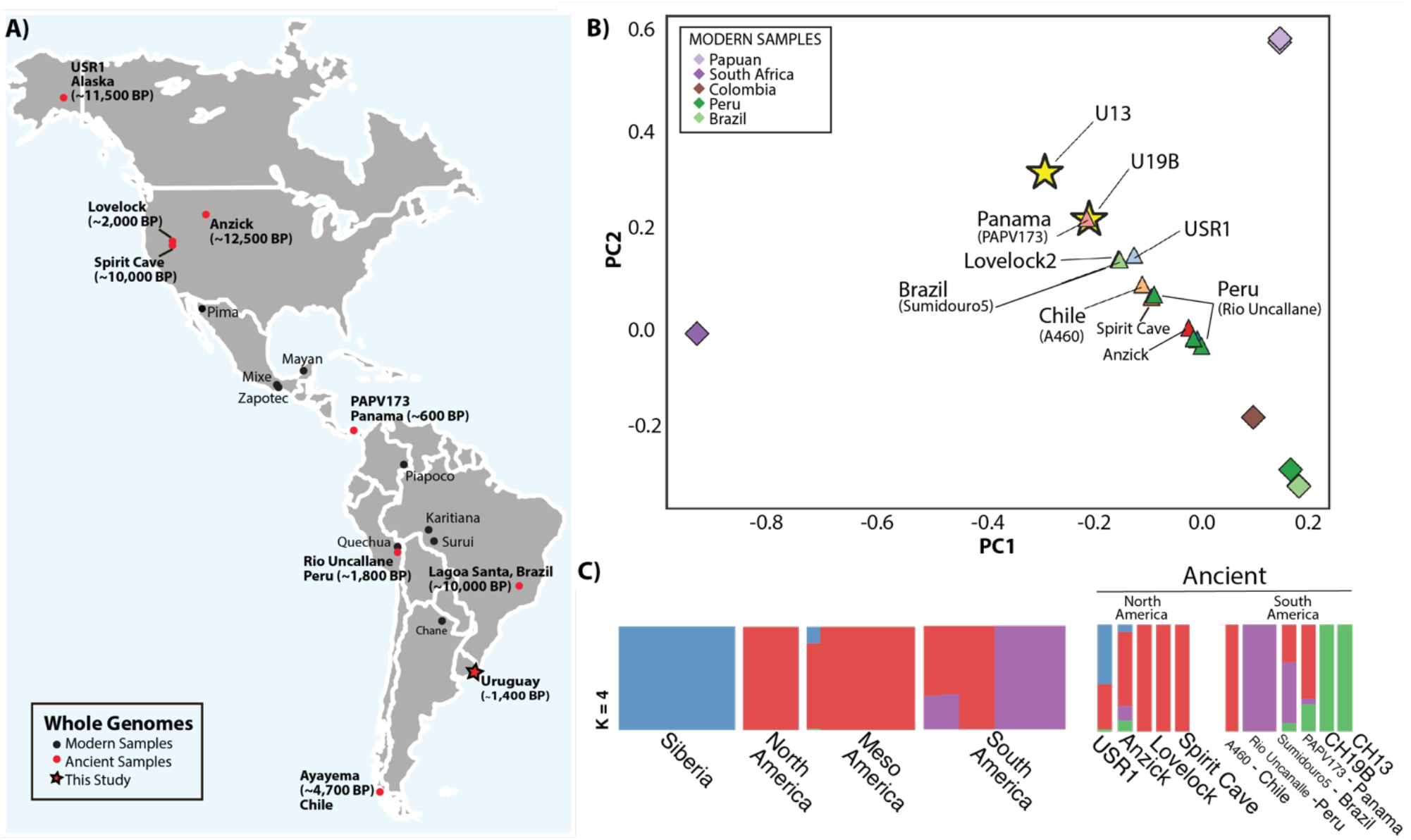
Assessing the genetic affinity of ancient Uruguay with the Americas. A) Map of ancient and modern whole genomes used in this study(3–8). B) Principal component analysis projection of the ancient Uruguay samples on to the first two principal components (PCs). C) Ancestry clusters generated with ADMIXTURE(25) of modern and ancient genomes from the Americas at *K*=4 clusters, which was chosen through cross-validation

### Mitochondrial DNA

The mitochondrial genome of CH-13 carries all diagnostic mutations of haplogroup C1d, namely 194T and 16051G, including the additional mutations 12378T, 16140C and 16288C which place it within subhaplogroup C1d3. This subhaplogroup had its origin approximately 9,000 years ago and apparently evolved entirely in what is now Uruguay, and is found also in CH-20, an individual found in the same site but between 700 and 1000 years older(10). The mtDNA of CH-19B carries mutations 1888A and 15930A, diagnostic of haplogroup C1c, but lacks further mutations associated with registered subhaplogroups (C1c1 through C1c8). Furthermore, CH-19B carries 606G and a deletion in position 7471, neither of which have been found in published mitochondrial sequences. The lineage represented by CH-19B might very well be extinct.

### Genomic analyses

We performed a principal component analysis to better understand the relationship of the ancient Uruguay individuals with other ancient individuals from the Americas. C/T and G/A transitions were removed from the dataset to guard against the most common forms of postmortem DNA damage(11) and to prevent false affinities among the ancient samples. Since the Uruguay ancient individuals are low coverage compared to the modern and ancient high coverage samples in the reference panel(3–6, 8, 12), we projected the ancient genomes onto top two principal components identified from the modern samples using SmartPCA(13). Interestingly, the contemporary ancient samples from Uruguay (CH19B, ∼1400 BP) and Panama (PAPV173(8), ∼1400 BP) show a strong affinity on the first two principal components (Fig. 1B), with the younger Uruguay sample (CH13, ∼600 BP) showing a more distant relationship. To further elucidate the relationship among the ancient Uruguay individuals and the Americas, we performed an ADMIXTURE-based cluster analysis, with *K*=4 clusters showing the best cross-validation value (Fig. 1C). Transitions were also removed for this analysis. The Uruguay ancient samples exhibit a green ancestry cluster which is shared with USR1(5) (Alaska, ∼11,500 BP) and Anzick-1(4) (Montana, ∼12,500 BP). In relation to the South America, the green cluster is shared with Sumidouro5(6) (Brazil, ∼10,000 BP) and PAP173(8) (Panama, ∼1400 BP), which shows the largest shared fraction. With regard to the modern samples in the reference panel, the green cluster is apparent in a Mayan individual(3) but is not observed in other populations.

To further assess the relationship of the ancient individuals from Uruguay with global populations, we utilized outgroup *f*3 statistics to assess the shared genetic ancestry with the modern individuals from Simons(3). Outgroup *f*3 statistics of the worldwide dataset demonstrate that both ancient Uruguay individuals display greater affinity with Indigenous groups from South America than with other populations (Figs. 2A and 2C). Ranked outgroup *f*3 statistics suggest that both ancient individuals from the two time periods (CH19B: ∼1450 BP and CH13: ∼668 BP) tend to share the greatest affinity with Brazilian living groups, the Surui and Karitiana (Fig. 2B). We also note that we do not see an Austronesian signal with either individual, which may suggest a more complicated ancestry despite the apparent affinity.

**Figure 2.**
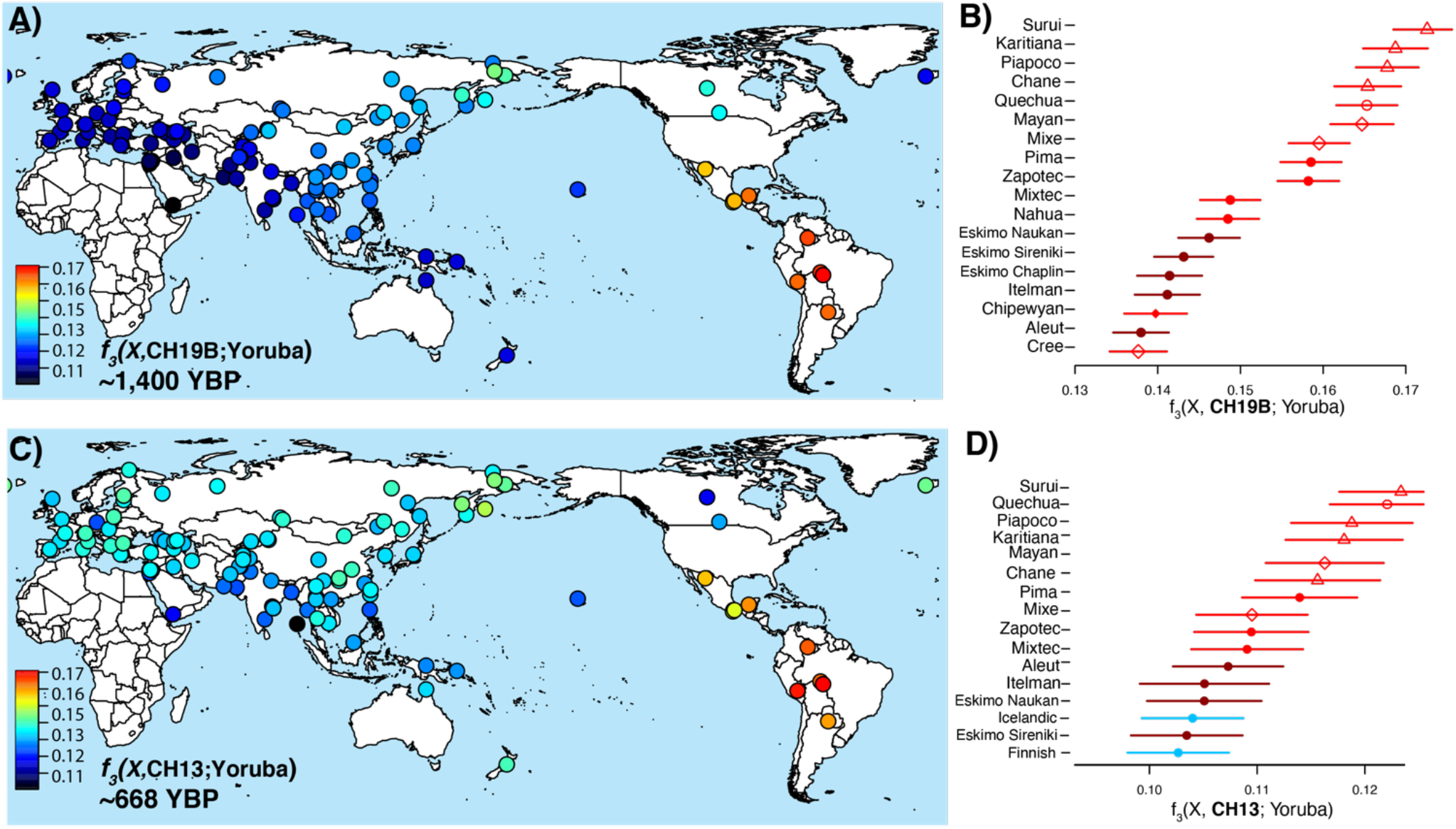
Outgroup *f*_3_ statistics. Left (A and C): Heat maps represent the outgroup *f*_3_ statistics quantifying the amount of shared genetic drift between the ancient Uruguay individuals and each of the contemporary populations from the Simons Genome Diversity Project^7^ since their divergence with the African Yoruban population. Right (B and D): Ranked *f*_3_ statistics showing the greatest affinity of the ancient Uruguay with respect to indigenous populations of the Americas.

To further explore the relationships between the ancient Uruguay individuals and the Americas, we utilized maximum likelihood trees inferred with TreeMix(14). Transitions were also removed from this analysis to prevent false affinities among ancient samples. Though migration events were not well supported by our bootstrapping validation, we do have support for the structure of a tree that shows a nuanced relationship between the ancient Uruguay individuals and South America. The ∼10,000 BP ancient sample from Brazil, Sumidouro5, falls basal to both ancient Panama and ancient Uruguay (Fig. 3A). It should be noted that Sumidoror5 is from an archeological site on the by the eastern coast of Brazil, as opposed to the Amazon populations, the Surui and Karaitiana, which are located in the western part of the country (Fig. 1A). Though we are careful to claim definitive relationships among the ancient and modern samples, the tree does seemingly correctly position the ancient individuals from the highlands of Peru (Rio Uncanalle(7)) with modern-day individuals from the same region, the Quechua(3). Furthermore, the ∼11,500-year-old USR1(5), a terminal individual from Alaska, is placed as an outgroup to all populations from the Americas, which lends additional validation of the tree structure.

**Figure 3.**
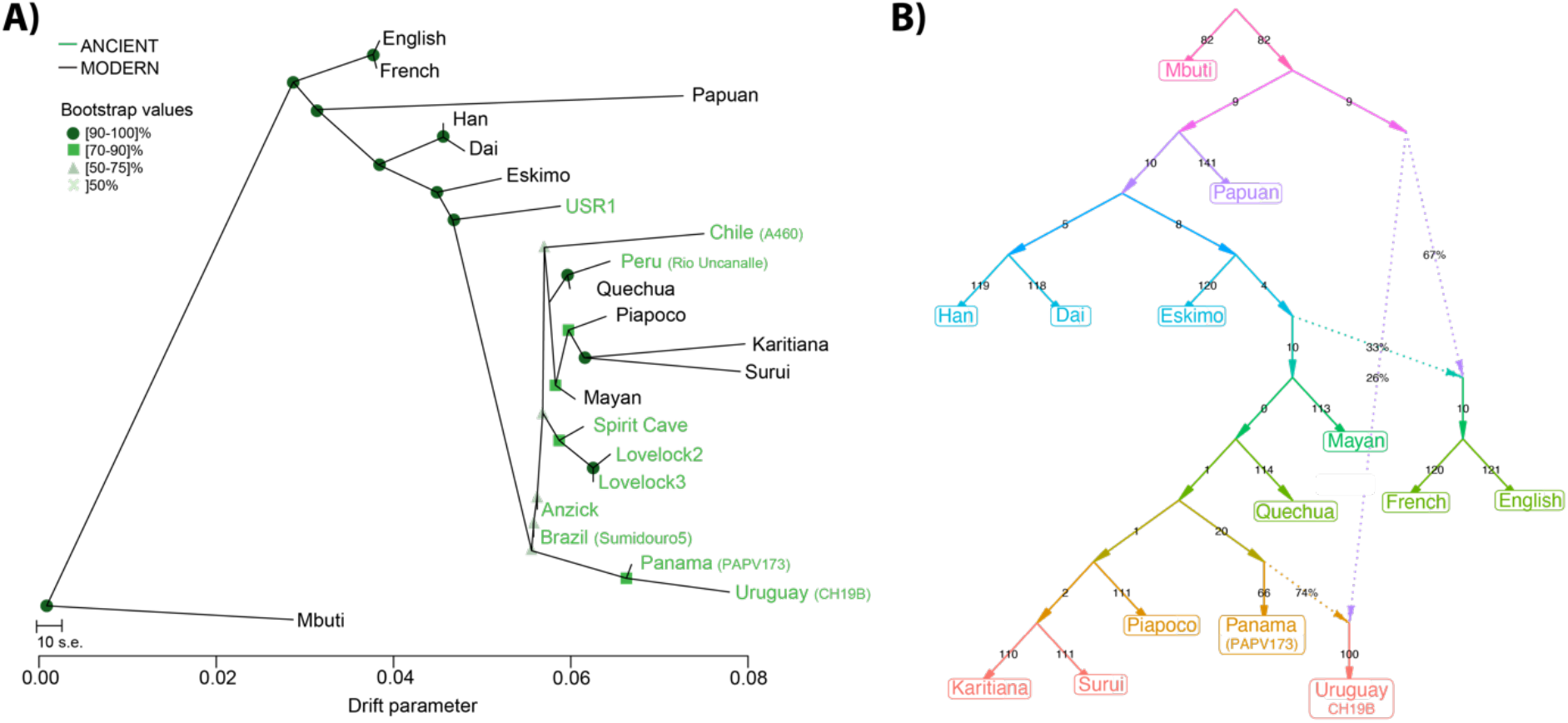
Maximum likelihood Tree and qpGraph. A) Maximum likelihood trees generated by TreeMix (14)using whole-genome sequencing data from the Simons Genome Diversity Project(3). The tree shows a connection between ancient samples from eastern Brazil, Panama, and Uruguay. B) The qpGraph best fitting model with two migration events. The populations included derive from the Simons(3) and shows a deep ancestral event in the direction of the ancient Uruguay sample (CH19B), along with a migration signal associated with ancient Panama.

We also tested the relationship of the 1,400-year-old ancient Uruguay individual with modern South American populations using qpGraph(15), which extends the *f*_3_ statistic to numerous populations by incorporating the topology of an admixture graph. We found that that the model fit best with two migration events, incorporating the individuals from key populations of the Simons Genome Diversity Project(3) (Fig. 3B). The topology of the graph also suggests that ancestry of the ancient Uruguay sample is deriving from two sources: a deep ancestral source and a source that led to the Karaitiana and Surui of Brazil. Both the maximum likelihood tree and the qpGraph show a more complicated picture than what was shown with the outgroup *f*_3_ statistic, whereby the Amazonian populations share a more distant connection to the ancient Uruguay individuals. This connection may relate to a more general ancestry signal from South America, rather than a direct one, and may be a consequence of different migrations upon entry into the continent. In contrast, we see a connection with ancient Brazil, Panama, and Uruguay on the maximum likelihood tree, where they form their own branch (Fig. 3A). The qpGraph shows a connection between the ancient Panama sample and the oldest Uruguay individual, demonstrating a migration event between the two (Fig. 4B). Taken together, it’s possible that the connections are reflective of migration routes that occurred along the Atlantic coast of South America. We also note a deep ancestral component contributing to the ancient sample form Uruguay (Fig. 3B), which combined with the ancestry cluster results (Fig. 1C), may represent a previously undetected ancestry in South America.

**Figure 4.**
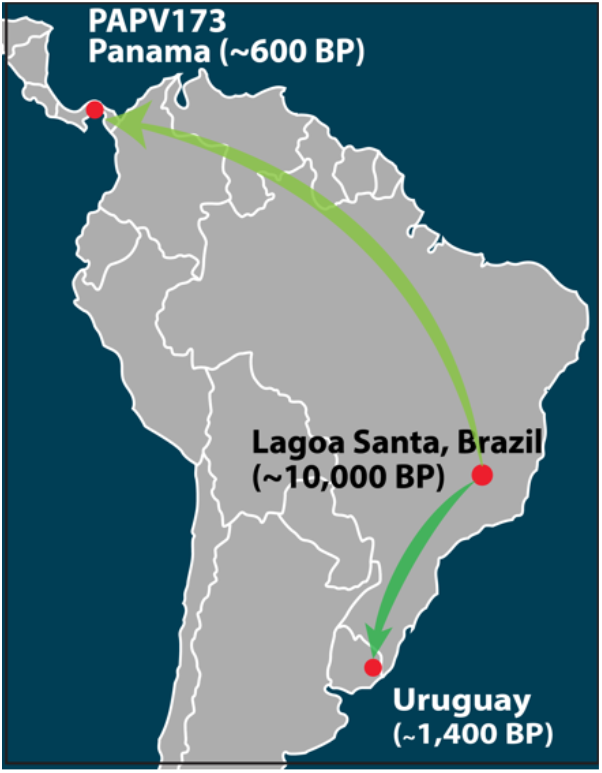
Atlantic coast migration route.

## Discussion

Here we start to elucidate the origins of the Indigenous people of Uruguay. We find that the ancestral population of the ancient individuals may have derived from a migration that stemmed closer to the Atlantic coast and may be different than the migrations that led to modern Amazonian indigenous populations from Brazil. We also find a close relationship to an ancient Panama sample, which could correlate with the source of the migration, Sumidouro Cave (part of the Lagoa Santa archaeological area), in Brazil, some 5,000 km away (Fig. 4).

That being said, besides indicating a past migration to Uruguay from Lagoa Santa through the Atlantic coast, this finding also suggests the existence of a putative south-to-north ancient migration route linking Lagoa Santa and Panama. Moreover, these movements seem to have occurred independently of the migratory waves that occurred near the Pacific coast in South American populations. Altogether, this presents genomic evidence for ancient migration events along, or at least closer to, South America’s Atlantic coast, which is supported by paleoenvironmental and chronological(16).

While we begin to unravel the relationship of the Indigenous people of Uruguay on a continental level, in addition to the potential discovery of a distinct ancestral component in South America, it also important to point out the need for ancient DNA from other archaeological sites from across Uruguay, especially those close to the time of European contact. Such samples will help to better capture the genetic diversity of the Indigenous that existed upon the Spanish in the 16^th^ Century, and by extension, better understand the diversity of Indigenous groups that existed. In doing so, future DNA studies could assist living people in Uruguay to potentially identify indigenous ancestry that is not limited to the “Charrúa.”(17).

## Supplemental Information

### Extended Methods

#### Archaeology and Samples

Site CH2D01 is a group of two mounds (A and B) located on the edge of the San Miguel wetlands, in the department of Rocha, in eastern Uruguay. The radiocarbon dating place the occupation of mound A between 2000 BP and the period of European contact. The mound is approximately 1.20 m high with a diameter of 35 m and is presumed to be an area of differentiated activity within a larger site of about 20,000 square meters. The archaeological materials recovered from mound A do not show clear spatial arrangements and were interpreted as “displaced primary contexts”: materials that were carried along with the sediments that make up the mound from the places where the activity was carried out. Implicit in this interpretation is the intentional character of the mound construction. Although there is ongoing debate about the exact mechanism of the formation of the “Cerritos,” there have been no subsequent interpretations about site CH2D01. In excavation IA, a 25 square meter excavation carried out in the center of mound A, several bone assemblages representing the primary and secondary burials of at least 21 individuals were recovered.

#### Ancient DNA and Sequencing

The ancient tooth samples were extracted, and sequencing libraries were constructed at the Lindo Ancient DNA laboratory at Emory University using the Dabney protocol(18). Libraries were prepared with the NEB Ultra II DNA Library Prep for Illumina, with modifications for ancient DNA, which including quartering the reagents, the utilization of 1:20 adaptor dilution, and 1.5 ul of premixed NEB indexes. Samples were preliminarily screened for endogenous DNA on the Illumina iSeq 100 at the Lindo Lab, with libraries that were not treated with the USER enzymes. Samples that were selected for deep sequencing on the NovaSeq 6000 at Dante Labs (L’Aquila, Italy), included libraries treated with the USER enzyme to help compensate for DNA damage.

The ancient raw sequences were trimmed for Illumina adapters using AdapterRemoval2(19) and aligned to the hg19 human reference sequence using the BWA mem algorithm(20), which has been shown to increase accuracy with ancient DNA mapping over the *aln* algorithm(21). The alignments from the shotgun sequences that were not treated with the USER enzyme were used to validate their ancient authenticity with MapDamage2(22). Both ancient individuals showed deamination patterns consistent with ancient DNA (Supp. Fig. 1).

Genotypes were called from both ancient individuals using the ancient DNA caller ARIADNA, which uses a machine learning method to overcome issue with DNA damage and contamination(9). The resulting VCF was further filtered to remove genotype calls with allele counts below 3. Since CH-19B showed a relatively high contamination rate, we further filtered the associated VCF using RFMix (23) to identify and remove sites that showed a high probability of deriving from Europeans. The VCFs were then merged with modern and ancient samples from the Americas with bcftools.

#### SmartPCA

We conducted Principal Component Analysis using the ‘smartpca’ program from the EIGENSOFT v7.2.1 package. Principal Components (PCs) were estimated with the ‘poplistname’ option and using representative individuals from present-day Native American and indigenous South African and Papuan populations from Simons Genome Diversity Panel(3). The ancient individuals were then projected onto the computed PCs with the ‘lsqproject: YES’ option. No outliers were excluded, and the analysis was based on 4,978,400 loci.

#### Assessment of population structure using ADMIXTURE

We started with the identical filtered dataset of called genotypes described above. We further pruned the dataset by removing sites in strong linkage disequilibrium (*r*^2^ > 0.1) using PLINK(24). The program *ADMIXTURE* was used to assess global ancestry of the ancient and present-day samples from this study. We computed cluster membership for *K*=2 through *K*=15 and found the lowest cross-validation value to be at *K=*4. The PONG program was used to visualize the admixture plots.

#### Outgroup f_3_ analysis

We applied the *qp3Pop* module of *ADMIXTOOLS*(15) to compute f3 statistics with the target population as the African Yoruban population and the two reference populations set as one of the ancient Uruguay samples (CH13 or CH19B) and the other as one of the non-African populations from the Simons Genome Diversity Project(3). For this analysis we retained C/T and G/A transitions, as the CH13 and CH19B ancient samples have been treated with uracil-DNA glycosylase to guard against this form of DNA damage.

#### *TreeMix* analysis

We started with the identical filtered dataset of called genotypes described above. *TreeMix* was applied the dataset to generate maximum likelihood trees and admixture graphs from allele frequency data. The Mbuti from the Simons dataset was used to root the tree (with the *–root* option). We accounted for linkage disequilibrium by grouping *M* adjacent sites (with the *–k* option), and we chose *M* such that a dataset with *L* sites will have approximately *L/M* ≈ 20,000 independent sites. At the end of the analysis (i.e., number of migrations) we performed a global rearrangement (with the *–global* option). We considered admixture scenarios with *m* = 0 and *m* = 3 migration events. Each migration scenario was run with 500bootstrapping replicates, and the replicateswere used to determine the confidence of each node.

#### *qpGraph* analysis

We employed the *ADMIXTOOLS2* (https://uqrmaie1.github.io/admixtools/index.html, *ADMIXTOOLS2* is currently under preparation) R package to perform *qpGraph*(15)estimation. We extracted *f*_2_ statistics between population pairs using a two megabase SNP block size while retaining C/T and G/A transitions, as the CH19B ancient sample has been treated with uracil-DNA glycosylase to guard against this form of DNA damage. We considered scenarios with *M* ∈ {0,1,2,3,4,5,6} migration events, with graph searches initiated by a random graph and Mbuti population set as the outgroup, stopping the search after 100 generations. If the best graph with *M* events did not have a better score than those with fewer events, then the graph search was rerun. The best-fit model of two migration events was chosen by assessing statistical differences between model score distributions computed from 500 bootstrap replicates of *f*_2_ blocks.

## Acknowledgements

This work was supported by National Science Foundation grants BCS-1926075, BCS-1945046, and BCS-2001063, and by Fundação de Amparo à Ciência e Tecnologia de Pernambuco grant BFP-0191-7.04/20.

---

**Supplementary Figure 1.**
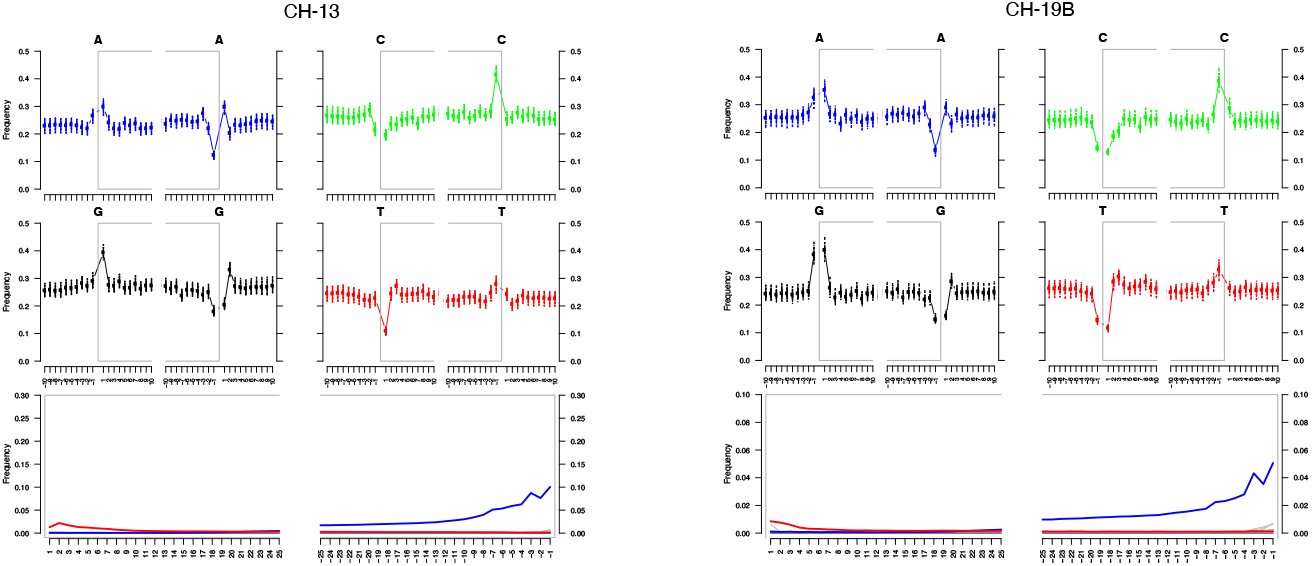
DNA Damage Patterns.

## References

1. Acosta y Lara, E.F., Los chaná-timbúes en la antigua Banda Oriental. Anales del Museo de Historia Natural de Montevideo VI, 1–27 (1955).

2. G. Renaud, V. Slon, A. T. Duggan, J. Kelso, Schmutzi: estimation of contamination and endogenous mitochondrial consensus calling for ancient DNA. Genome Biol. 16, 47 (2015).

3. S. Mallick, et al., The Simons Genome Diversity Project: 300 genomes from 142 diverse populations. Nature 538, 201–206 (2016).

4. M. Rasmussen, et al., The genome of a Late Pleistocene human from a Clovis burial site in western Montana. Nature 506, 225–229 (2014).

5. J. V. Moreno-Mayar, et al., Terminal Pleistocene Alaskan genome reveals first founding population of Native Americans. Nature 488, 370–9 (2018).

6. J. V. Moreno-Mayar, et al., Early human dispersals within the Americas. Science 525, eaav2621 (2018).

7. J. Lindo, et al., The genetic prehistory of the Andean highlands 7000 years BP though European contact. Science Advances 4, eaau4921 (2018).

8. M. R. Capodiferro, et al., Archaeogenomic distinctiveness of the Isthmo-Colombian area. Cell 184, 1706-1723.e24 (2021).

9. J. K. Kawash, S. D. Smith, S. K. D. Research, 2018, ARIADNA: machine learning method for ancient DNA variant discovery | DNA Research | Oxford Academic. academic.oup.com https://doi.org/10.1093/dnares/dsy029.

10. M. Sans, et al., A South American Prehistoric Mitogenome: Context, Continuity, and the Origin of Haplogroup C1d. PloS one 10, e0141808 (2015).

11. A. W. Briggs, et al., Patterns of damage in genomic DNA sequences from a Neandertal. PNAS 104, 14616–14621 (2007).

12. J. Lindo, et al., Ancient individuals from the North American Northwest Coast reveal 10,000 years of regional genetic continuity. Proc Natl Acad Sci USA 114, 4093–4098 (2017).

13. N. Patterson, A. L. Price, D. Reich, Population Structure and Eigenanalysis. Plos Genet 2, e190 (2006).

14. J. K. Pickrell, J. K. Pritchard, Inference of Population Splits and Mixtures from Genome-Wide Allele Frequency Data. PLoS Genet 8, e1002967 (2012).

15. N. Patterson, et al., Ancient Admixture in Human History. Genetics 192, 1065–1093 (2012).

16. Miotti, “La fachada atlántica, como puerta de ingreso alternativa de la colonización humana de América del Sur durante la transición Pleistoceno/Holoceno.” in Simposio Internacional Del Hombre Temprano En América, J. C. J. López, Ed. (Instituto Nacional de Antropología e Historia., 2006), pp. 155–188.

17. L. Spangenberg, et al., Indigenous Ancestry and Admixture in the Uruguayan Population. Frontiers Genetics 12, 733195 (2021).

18. J. Dabney, et al., Complete mitochondrial genome sequence of a Middle Pleistocene cave bear reconstructed from ultrashort DNA fragments. Proc Natl Acad Sci USA 110, 15758–15763 (2013).

19. M. Schubert, S. Lindgreen, L. Orlando, AdapterRemoval v2: rapid adapter trimming, identification, and read merging. BMC Res Notes 9, 395 (2016).

20. H. Li, R. Durbin, Fast and accurate short read alignment with Burrows–Wheeler transform. Bioinformatics 25, 1754–1760 (2009).

21. W. Xu, et al., An efficient pipeline for ancient DNA mapping and recovery of endogenous ancient DNA from whole-genome sequencing data. Ecol. Evol. 11, 390–401 (2020).

22. H. Jónsson, A. Ginolhac, M. Schubert, P. L. F. Johnson, L. Orlando, mapDamage2.0: fast approximate Bayesian estimates of ancient DNA damage parameters. Bioinformatics 29, 1682– 1684 (2013).

23. B. K. Maples, S. Gravel, E. E. Kenny, C. D. Bustamante, RFMix: A Discriminative Modeling Approach for Rapid and Robust Local-Ancestry Inference. The American Journal of Human Genetics 93, 278–288 (2013).

24. C. C. Chang, et al., Second-generation PLINK: rising to the challenge of larger and richer datasets. Gigascience 4, 7 (2015).

25. M. Harris, M. DeGiorgio, Admixture and Ancestry Inference from Ancient and Modern Samples through Measures of Population Genetic Drift. Hum. Biol. 89, 21–46 (2017).

